# Broad spectrum antibiotic-degrading metallo-β-lactamases are phylogenetically diverse and widespread in the environment

**DOI:** 10.1101/737403

**Authors:** Marcelo Monteiro Pedroso, David W. Waite, Okke Melse, Liam Wilson, Nataša Mitić, Ross P. McGeary, Iris Antes, Luke W. Guddat, Philip Hugenholtz, Gerhard Schenk

## Abstract

Antibiotic resistance has emerged as a major global health threat. The Zn^2+^-dependent metallo-β-lactamases (MBLs) are of particular concern as they act on the most widely prescribed class of antibiotics, the β-lactams, and are largely unaffected by commonly used β-lactamase antagonists such as clavulanic acid. MBLs are subdivided into three groups (B1 to B3); despite low overall sequence similarity, their catalytic centers are conserved with two closely spaced Zn^2+^ binding sites (α and β site). We recovered almost 1500 B3 MBLs from >100,000 public microbial genomes representing a wide range of habitats including pristine sites not impacted by human activity. Although homologs were predominantly identified in members of the bacterial phylum *Proteobacteria*, the recovered B3 MBLs represent a much broader phylogenetic diversity than is currently appreciated based on the study of model pathogens. This includes three active site variants inferred to have arisen from the ancestral B3 enzyme. One of these variants, B3-RQK, is noteworthy for being broadly sensitive to clavulanic acid. Through targeted mutations we demonstrate that the presence of a lysine residue (Lys263) in the β site of the catalytic center of this variant confers sensitivity to this compound. Replacing this lysine with the canonical histidine (His263) found in all other MBLs restored resistance. Crystallographic and computational data reveal that clavulanic acid inhibits B3-RQK MBLs by displacing the Zn^2+^ ion in the β site. Therefore, modifying clavulanic acid to effectively interact with His263 may increase the therapeutic range of this widely used antibiotic resistance drug.

**Significance:** This study surveys the environmental and phylogenetic diversity of the B3 subgroup of antibiotic-degrading metallo-β-lactamases (MBLs). B3-like MBLs are more widespread in the environment than previously appreciated suggesting multiple unrecognized reservoirs of antibiotic resistance. Three variants of the canonical active site were identified, including B3-RQK, which amongst the B3 MBLs is uniquely inhibited by the antibiotic resistance drug clavulanic acid. We demonstrate that the mode of inhibition involves the displacement of a catalytically essential Zn^2+^ ion from the active site. It may thus be possible to modify clavulanic acid so that it can compete with the Zn^2+^ ions in other MBLs as well, increasing the therapeutic range of this compound.

---

Antibiotic resistance has emerged as a major threat to global health (1, 2). Multi-drug resistant bacteria already kill more patients in the United States each year than HIV/AIDS, Parkinson’s disease, emphysema and homicide combined (3-5). Among the most effective bacterial resistance mechanisms are β-lactamases, which are enzymes that inactivate many of the commonly used β-lactam-based antibiotics (*e*.*g*. penicillin) (1, 6). β-Lactamases are divided into four distinct classes: Classes A, C and D (the serine-β-lactamases, SBLs) use an active site serine residue to initiate cleavage of the antibiotic, and Class B (the metallo-β-lactamases, MBLs) use a metal ion (usually Zn^2+^)-activated hydroxide to promote cleavage (1, 7). Clinically relevant inhibitors of Class A and D SBLs are available and in use (*e*.*g*. clavulanic acid), but for MBLs the search for such inhibitors has remained challenging (8). Consequently, pathogens carrying MBL genes pose a major health risk.

Conventionally, MBLs have been divided into three subgroups, *i*.*e*. B1, B2 and B3 (9). Enzymes of the B1 subgroup constitute the majority of MBLs associated with antibiotic resistance; members include IMP-1, NDM-1, VIM-1 and various derivatives (mutants) thereof (10-12). For optimal catalytic efficiency these enzymes require two closely spaced Zn^2+^ ions, located in the α- and β-sites of the catalytic center. These two sites are characterized by the sequence motif His116, His118, His196 in the α-site and Asp120, Cys221, His263 in the β-site (*i*.*e*. HHH/DCH) (13). Fewer B2-type MBLs are currently known but they are characterized by a strong preference for “last line” carbapenem substrates such as meropenem (14). Despite being homologous and sharing a high degree of sequence similarity with the B1-type MBLs in the active site (*i*.*e*. NHH/DCH for the α/β site motifs), members of the B2 subgroup only require one Zn^2+^ for catalysis (located in the β-site); the presence of a second metal ion leads to inhibition (1, 7, 15). Only a small number of B3-type MBLs have been identified, including L1 (16-20), FEZ-1 (21, 22), AIM-1 (23-25), GOB-1 (26, 27), LRA-8 (28), MIM-1 and MIM-2 (29, 30). They have a structurally similar active site motif to B1 MBLs (HHH/DHH), and also require a bimetallic Zn^2+^ center for optimal catalytic efficiency (1, 7, 13, 31). B3 MBLs, however, have low sequence similarity to B1 and other MBLs (<20% amino acid identity) instead being more closely related to Class D SBLs (>30% amino acid identity) (32).

Recently, SPR-1, a B3 MBL with an atypical active site (His116, Arg118, His196 in the α site and Asp120, Gln121, Lys263 in the β site, *i*.*e*. H**R**H/D**QK**) was identified in *Serratia proteamaculans* via its sequence similarity to AIM-1 (33). While the observed active site substitutions (in bold) do not significantly alter the substrate specificity, they affect the enzyme-metal ion interactions in the active site. Only one Zn^2+^ binds in the resting state and the enzyme is activated by the substrate-promoted interaction with a second metal ion (33). Such an activation mechanism, while observed in other metalloenzymes (34, 35), is not common amongst MBLs (36). It was thus proposed that this B3 variant may represent a distinct subgroup of MBLs (33, 37), for which we propose the abbreviated name B3-RQK based on its active site substitutions (*see above*). Although not recognized in the study, another B3-RQK MBL (CSR-1) was subsequently identified in an invasive foodborne pathogen, *Cronobacter sakazakii* (38). This organism can cause life-threatening meningitis, septicemia and necrotizing enterocolitis with a 40-80% mortality rate (38-40), and resistance to antibiotics amongst various *Cronobacter* strains has already been reported (38). Another B3 MBL (GOB-1) with a minor motif variant (**Q**HH/DHH; B3-Q) was discovered in the opportunistic pathogen *Elizabethkingia meningoseptica*, which has broad spectrum activity against a range of β-lactam antibiotics, similar to native B3 MBLs such as AIM-1 (26, 27).

The discovery of MBLs with atypical active site sequences indicates that these enzymes and associated antibiotic-inactivating mechanisms are more diverse than previously appreciated. This could have important implications for the design of clinically useful broad-spectrum inhibitors, which most often are designed to target a conserved active site. In order to gain a deeper understanding of B3 MBLs, we recovered almost 1500 homologs from a large public dataset of microbial genomes representing a wide range of habitats. Phylogenetic inference indicated that the B3 MBL dataset greatly increases the known diversity of this gene family, including identification of three active site motif variants (B3-RQK, B3-Q, and B3-E). Among these variants B3-RQK is notable as it has a number of unusual properties including an occluded active site in the mature protein, reduced metal binding and catalytic activity, and reduced ability to confer *ex vivo* resistance. Furthermore, B3-RQK enzymes are inhibited by clavulanic acid. Through targeted mutations, we pinpointed the cause of this inhibition to substitution of the canonical histidine to a lysine in position 263 of the β site in the catalytic center, which could provide a focus for drug development.

## Results and Discussion

### Phylogenetic analysis of B3 MBL homologs

Release 02-RS83 of the Genome Taxonomy Database (41) comprising 111,330 quality-filtered bacterial and archaeal genomes was searched for homologs of B3 MBLs in order to assess the evolutionary history and diversity of this family of antibiotic-degrading enzymes. A total of 1,449 putative B3 MBL proteins were identified in 1,383 genomes (representing 1.2% of all analyzed genomes), of which 1,150 have the canonical B3 active site residues (HHH/DHH) and 47 have the B3-RQK motif with changes in both the α- and β-sites (H**R**H/D**QK**). Two additional minor variants of the B3 motif with single amino acid changes in their α-sites were also discovered; the **Q**HH/DHH motif (B3-Q) previously noted in GOB-1 (26, 27) was present in 162 proteins, while the **E**HH/DHH motif (B3-E), identified in 90 proteins, has not previously been reported. Phylogenetic inference of a representative subset of 673 of these proteins indicates that each of the three motif variants originate from within the B3 radiation (**Figure 1**). B3-RQK appears to have only arisen once, likely because the ancestral change required at least four nucleotide substitutions to produce the three amino acid changes (**Figure 1**). By contrast, the B3-Q and B3-E variants have a single amino acid difference in position 116 requiring only one and two nucleotide changes, respectively. The B3-Q variant appears to have arisen on at least five independent occasions and reverted back to the B3 motif on at least six occasions as a result of the need for only one nucleotide change (**Figure 1**).

**Figure 1.**
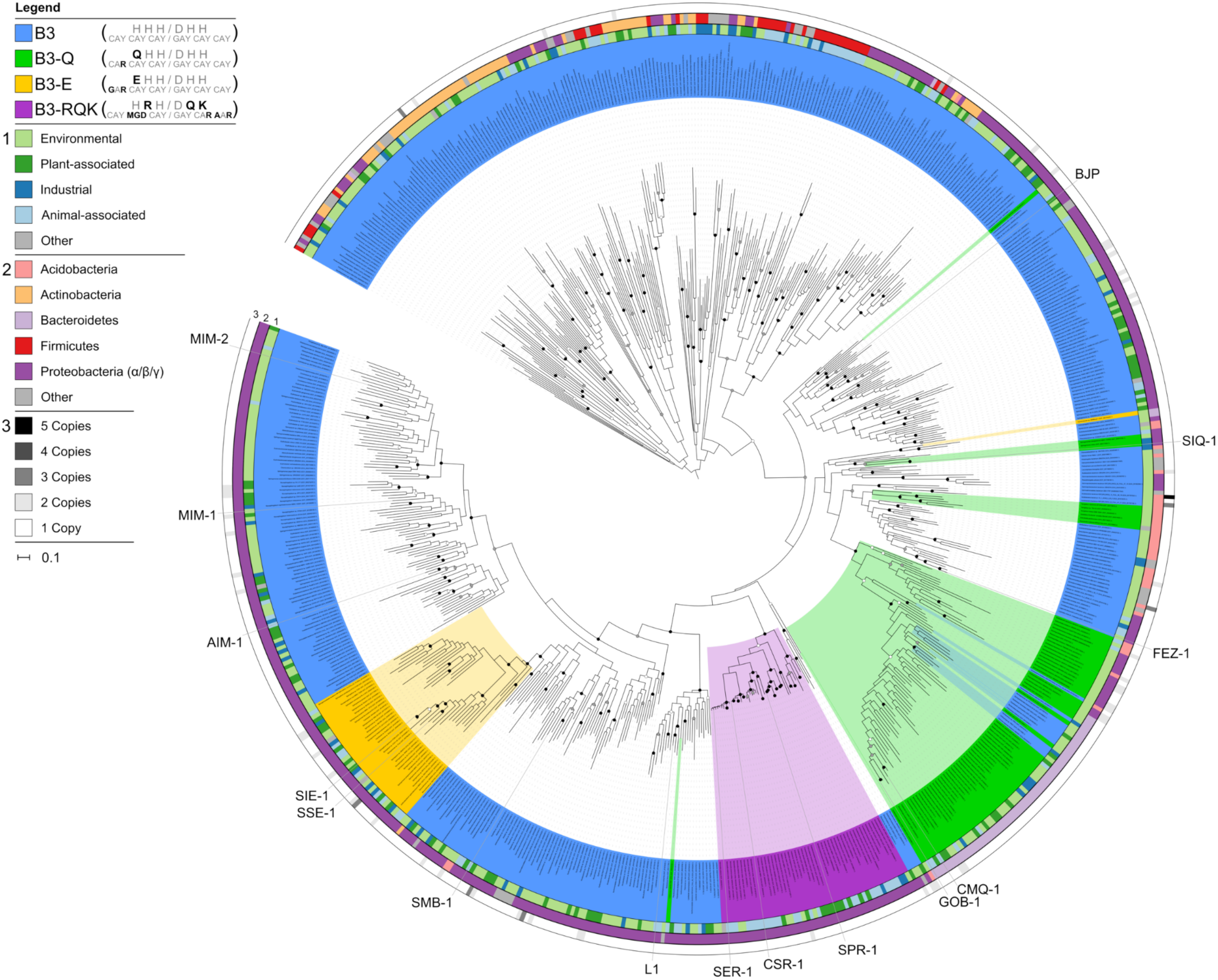
Unrooted maximum likelihood tree of MBLs belonging to subgroup B3, highlighting three active site variants. The tree was inferred from 673 dereplicated B3 MBLs identified in 1,383 bacterial genomes screened from a total of 111,330 bacterial and archaeal genomes. Bootstrap support for the interior nodes is indicated by filled (black: >90%, gray: >80%) or open (>70%) circles. B3 active site variants are indicated by different colors according to the legend in the top left of the figure. The inner circle (1) represents the phylum-level affiliations of the B3-containing bacteria. The middle circle (2) represents the habitat source of the B3-containing bacteria, and the outer circle (3) represents B3 gene copy number in each genome.

### Host taxonomy and environmental distribution

No archaeal genomes harbored B3 MBLs, and the majority were found in just four bacterial phyla; the *Proteobacteria, Actinobacteria, Bacteroidetes* and *Firmicutes* (**Figures 1 & 2**). While this certainly reflects to some extent the current over-representation of these phyla in the genome database (**Supplemental Figure 1**), it also suggests that the host range of B3 MBLs is relatively restricted. Overlaying the B3 MBL tree with the host phylogeny (ring 1 in **Figure 1**) indicates extensive lateral transfer of the MBL genes, most conspicuously at the phylum level. The B3-RQK and B3-E variants were found exclusively in the *Proteobacteria* and B3-Q were mostly identified in the *Bacteroidetes* (**Figure 2**). Localization of the B3-RQK genes in the family *Enterobacteriaceae* and genus *Acinetobacter* is particularly striking, suggesting that these changes to the active site are relatively recent and lateral transfer has been limited thus far. Between two and five B3 genes were found in 57 genomes, with the most copies being present in an as-yet-uncultured member of the *Acidobacteria* (**Supplemental Table 1**). Numerous instances of native B3 enzymes co-occurring with B3-E and B3-Q were identified, however, only one instance of a B3 and B3-RQK was found (in a member of the *Enterobacteriaceae*) possibly indicating functional incompatibility of these enzymes (**Supplemental Table 1**).

**Figure 2.**
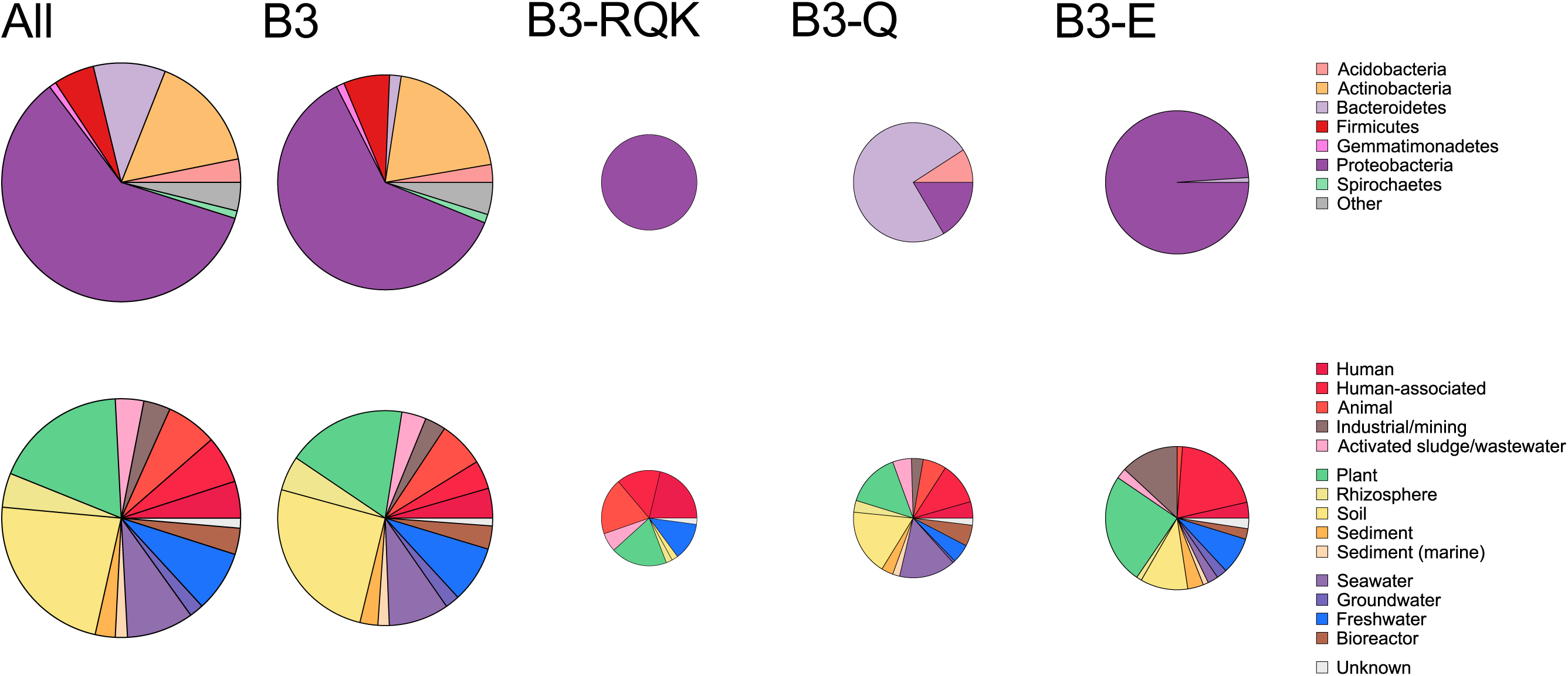
Bacterial host affiliation and environmental distribution of B3 enzymes. Pie charts are scaled by the number of component protein sequences in each enzyme category. Host affiliation is shown for the top seven phyla and environments are categorized according to the legend to the right of the figure.

A majority of habitats linked to the identified sequences are associated with humans (typically hospital or farming environments) or soil and rhizosphere samples, however there is a sizable environmental reservoir beyond these habitats (outer ring in **Figure 1** and **Figure 2**). Previously studied, clinically relevant MBLs were not clustered together as a single monophyletic unit, but rather interspersed throughout the tree. This finding suggests that clinical MBLs have entered into the hospital environment through multiple, independent incursion events and are not the result of a single human-associated MBL undergoing diversification. Notably, many B3-like MBLs were identified in water-borne bacteria which may be linked to human activity whereby human-associated MBLs are mobilized into water columns, and/or reflects aquatic habitats simply being under-appreciated reservoirs of B3 MBLs.

### B3 active site variant MBLs confer resistance to β-lactam antibiotics

The phylogenetic analysis of the B3 MBL family, including the three active site variants, indicates a large and diverse reservoir of bacterial species potentially able to degrade β-lactam antibiotics. Canonical B3 MBLs such as L1 and AIM-1 (**Figure 1**) are potent β-lactamases as shown by *in vitro* β-lactam antibiotic degradation assays and their ability to confer *ex vivo* resistance to *Escherichia coli* (24, 42-44). The only characterized representatives of B3-RQK (SPR-1 (33)) and B3-Q (GOB-1 and its variant GOB-18 (26, 27)) have β-lactamase activity, but only the latter have been shown to confer resistance to a host organism (45). We were therefore interested to determine if the ability to confer resistance to a bacterial host is a universal property among B3 enzymes.

Twelve genes representing each of the B3 active site motif variants were selected for plasmid expression of the corresponding mature proteins in *E. coli* BL21. Their ability to confer resistance against a range of substrates that represent the three major classes of β-lactam antibiotics was assessed using disc tests summarized using an aggregate resistance score (**Table 1 and Supplemental Table 2**). Characterized canonical B3 MBLs (L1, AIM-1, MIM-1, MIM-2) were included in the analysis as positive controls and plasmid-free BL21 strain used as a negative control. Aggregate resistance scores of the MBLs were calculated as the sum of individual resistance scores for each of nine substrates. The scoring system was 1 for resistance (R; zone of inhibition <18 mm), 0.5 for marginal resistance (r; inhibition zone 18-26 mm) and 0 for sensitivity (S; inhibition zone ≥27 mm) based on zones observed around the negative control. B3, B3-E and B3-Q MBLs conferred resistance to at least one antibiotic of each class, and based on their aggregate resistance scores the subgroups can be ranked from most to least resistant as follows: B3 > B3-E > B3-Q. The tested representatives of B3-Q and B3-E have similar resistance profiles, being completely sensitive only to meropenem and biapenem. In contrast, B3-RQK enzymes did not confer resistance to any antibiotic in *E. coli*. However, based on the observation that the B3-RQK enzyme SPR-1 was catalytically active only when it was further truncated at the N-terminus (49 amino acids (33)), we tested truncated versions of the mature proteins of all three B3-RQK representatives (*i*.*e*. SPR-1_trunc_, CSR-1_trunc_ and SER-1_trunc_; 32 to 49 N-terminal amino acids removed). The truncated enzymes could indeed confer resistance to *E. coli*, although all were susceptible to carbenicillin, cefuroxime and meropenem (**Table 1**). In summary, the modified active site enzymes had overall reduced ability to confer resistance to a bacterial host, but are still active against a broad range of substrates including last-resort antibiotics.

**Table 1.**
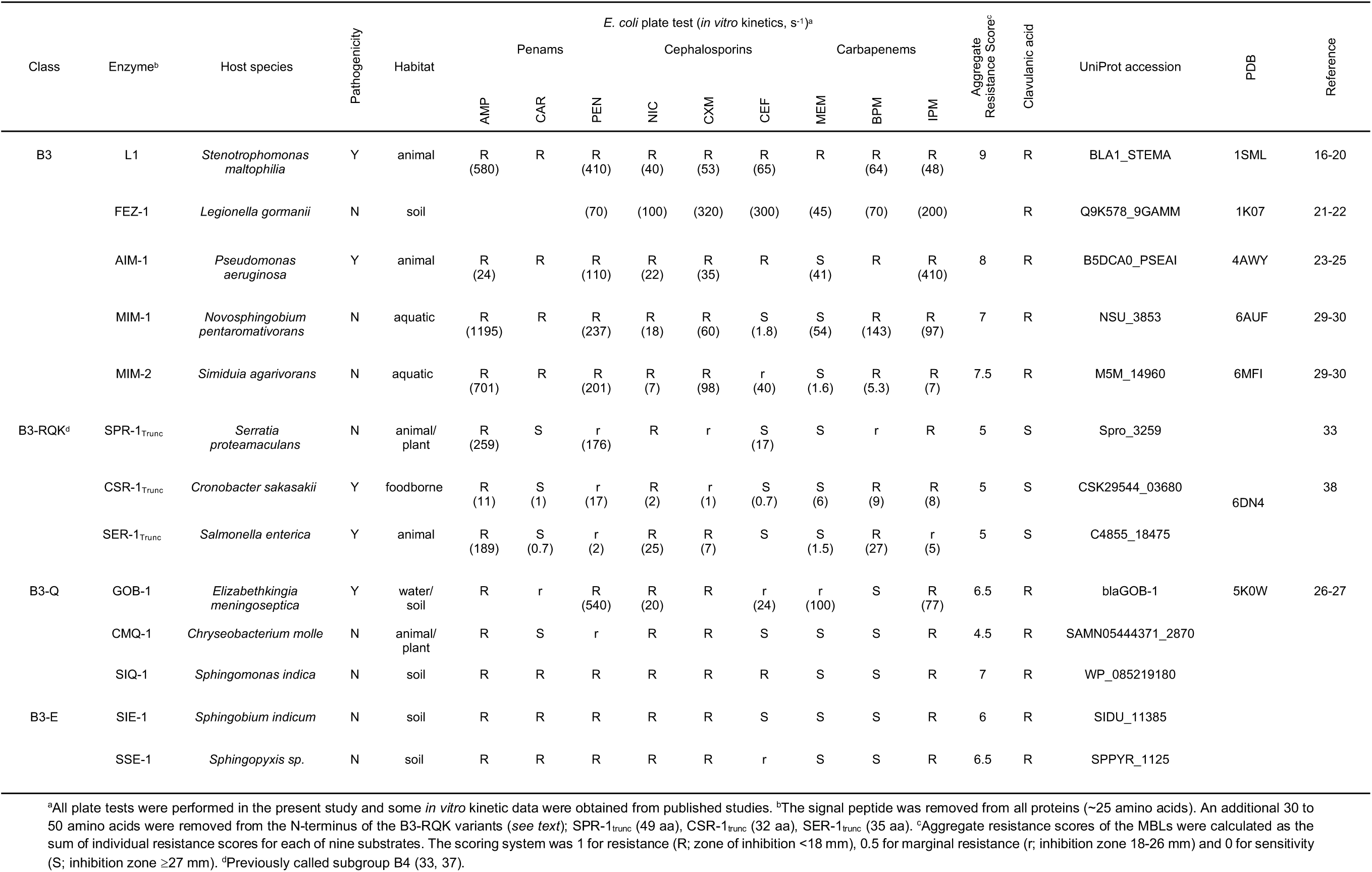
*In vitro* kinetic data and *ex vivo* plate tests of selected B3 metallo-β-lactamases.

### B3-RQK enzymes are sensitive to clavulanic acid

The most widely used clinical inhibitor of SBLs, clavulanic acid, is ineffective against the great majority of MBLs confounding efforts to inhibit their activity against antibiotics (2, 46). We tested the B3 variants for their resistance to clavulanic acid using a modification of the whole cell plate assay described above, whereby the inhibitor was added to the disc together with one of the tested antibiotics. Canonical B3 enzymes were resistant as were variants B3-E and B3-Q. However, the active, N-terminally truncated versions of the B3-RQK enzymes are sensitive to clavulanic acid (**Table 1**). The effect of clavulanic acid on the activity of expressed and purified B3 enzymes was tested using a standard *in vitro* kinetic assay. As expected, this inhibitor had no significant effect on the activity of B3, B3-E or B3-Q enzymes up to a concentration of 1 mM. However, CSR-1_trunc_, SPR-1_trunc_ and SER-1_trunc_ were inhibited by clavulanic acid even at concentrations as low as 10 µM (**Supplemental Figure 2**). The inhibitory effect, based on the constant *K*_*i*_, is 350 µM for CSR-1_trunc_, 201 µM for SPR-1_trunc_ and 290 µM for SER-1_trunc_, comparable to *K*_*i*_ values observed in SBLs (20 to 200 µM) (47), and in the only MBL for which a *K*_*i*_ for clavulanic acid (67 µM) has been reported (the B1 MBL SPM-1) (48).

### Structural and functional characterization of the B3-RQK variant

Among the motif variants, B3-RQK stands out due to the extent of changes to the active site, uniquely including two amino acid substitutions in the *β*-site (**Figure 1**), the requirement for N-terminus truncation of the mature protein to confer antibiotic resistance, and sensitivity to clavulanic acid. It was therefore of interest to structurally and functionally characterize this variant. We began by testing the enzymatic activity of CSR-1 and its homolog from *Salmonella enterica*, SER-1 (**Table 1**). As was noted for the first catalytically characterized B3-RQK representative (SPR-1), the mature proteins were not active until the N-terminus was truncated (33) (**Tables 1** and **2** and **Supplemental Table 3**). This led us to speculate that the N-terminus was occluding the substrate from binding to the active site. To test this hypothesis, we determined the crystal structure of the mature CSR-1 protein and CSR-1_trunc_, and compared them to available structures of canonical B3 (L1, AIM-1) and B3-Q (GOB-1) enzymes (**Figure 3**). A conspicuous difference between the four enzymes is the relative position of their N-termini. In L1, the N-terminus protrudes from the structure thereby making the active site accessible (16), whereas in AIM-1, a disulphide bridge between Cys32 and Cys66 locks its N-terminus in a position away from the catalytic center (**Figure 3**) (23, 25). Similar to L1, GOB-1 (**Figure 3**) has a protruding N-terminus exposing the active site (26). Only in the case of the mature CSR-1 structure is the N-terminal loop located above the active site, likely blocking substrate access. Removal of the N-terminus thus exposes the catalytic core (**Figure 3**).

**Figure 3.**
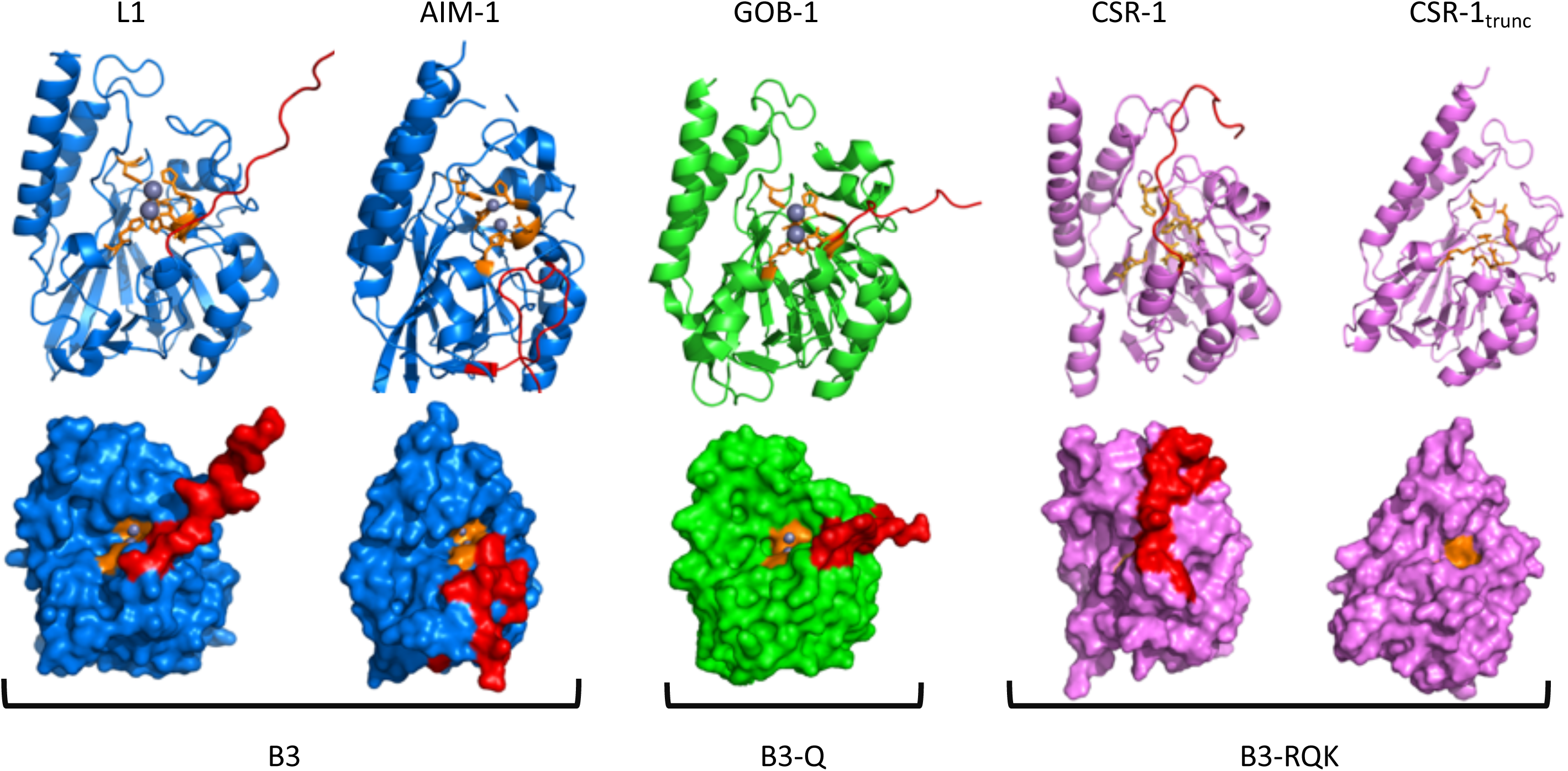
Selected crystal structures of native B3-MBLs and two active site variants. The names of the specific enzymes are indicated at the top of the figure (highlighted in **Figure 1**), and the type of active site is noted at the base of the figure. The top panel shows cartoon representations of the overall structure of the selected enzymes, with active site ligands in orange, Zn^+2^ ions in gray, and the N-terminal loop in red. The bottom panel shows surface representations of the five structures highlighting occlusion of the active site by the N-terminus in CSR-1.

The second unusual feature separating B3-RQK enzymes from canonical B3s and other variants is the composition of the active site (**Figure 1**). In all known MBL structures the cavity of the catalytic center can accommodate up to two Zn^2+^ ions (bound in the α- and β-site, respectively) (1, 7). However, no metal ions were observed in the crystal structures of CSR-1 and CSR-1_trunc_ (**Figure 3**). This absence is consistent with reduced metal ion affinity, which is likely due to the three distinct amino acid variations in the H**R**H/D**QK** active site sequence motif of B3-RQK MBLs (**Figure 1**). To probe the role of these three residues for metal binding, single (sm), double (dm) and triple (tm) mutants of CSR-1_trunc_ were generated. In CSR-1_trunc,sm_ R118 and in CSR-1_trunc,dm_ Q121 and K263 were replaced by histidines to create α or *β*-sites identical to those of canonical B3 MBLs, respectively (*i*.*e*. H**H**H or D**HH**). In CSR-1_trunc,tm_ R118, Q121 and K263 were all substituted by histidines, thus reconstituting the canonical B3 active site (*i*.*e*. H**H**H/D**HH**). These mutations had no significant effect on the overall structures (**Figure 3**), however, in CSR-1_trunc,sm_ electron density indicates the presence of one bound Zn^2+^ in the α-site, while in the double and triple mutants both the α- and β-sites are occupied by Zn^2+^ ions (**Figure 4a**). A third Zn^2+^ ion was observed to bind in the double mutant outside of the α- and β-sites, which we predict is a crystallographic artefact as it would block the active site. The qualitative observation that the three mutations significantly enhance metal binding was confirmed using isothermal titration calorimetry (ITC; **Supplemental Figure 3** and **Supplemental Table 4**). CSR-1 and CSR-1_trunc_ bind two Zn^2+^ ions with moderate to weak affinities (α*K*_*d*_ ∼2 µM and β*K*_*d*_ ∼150 µM), additionally indicating that the N-terminus of the mature enzyme does not affect metal ion binding. Two Zn^2+^ ions also bind to CSR-1_trunc,sm_, but the affinity of the α-site (α*K*_*d*_ ∼0.1 µM) is significantly enhanced, while the β-site (β*K*_*d*_ ∼180 µM) is unaffected by the mutation. In the double and triple mutants both the α and β metal binding sites have high affinity (α*K*_*d*_ = β*K*_*d*_ ∼0.14 µM and α*K*_*d*_ ∼0.06 µM and β*K*_*d*_ ∼0.4 µM, respectively). For comparison, similar ITC measurements with AIM-1 result in an α*K*_*d*_ and β*K*_*d*_ of ∼0.17 µM (24) indicating that the double and triple mutants of CSR-1 fully restore the characteristic ability of canonical B3 enzymes to bind two Zn^2+^ ions tightly.

**Figure 4.**
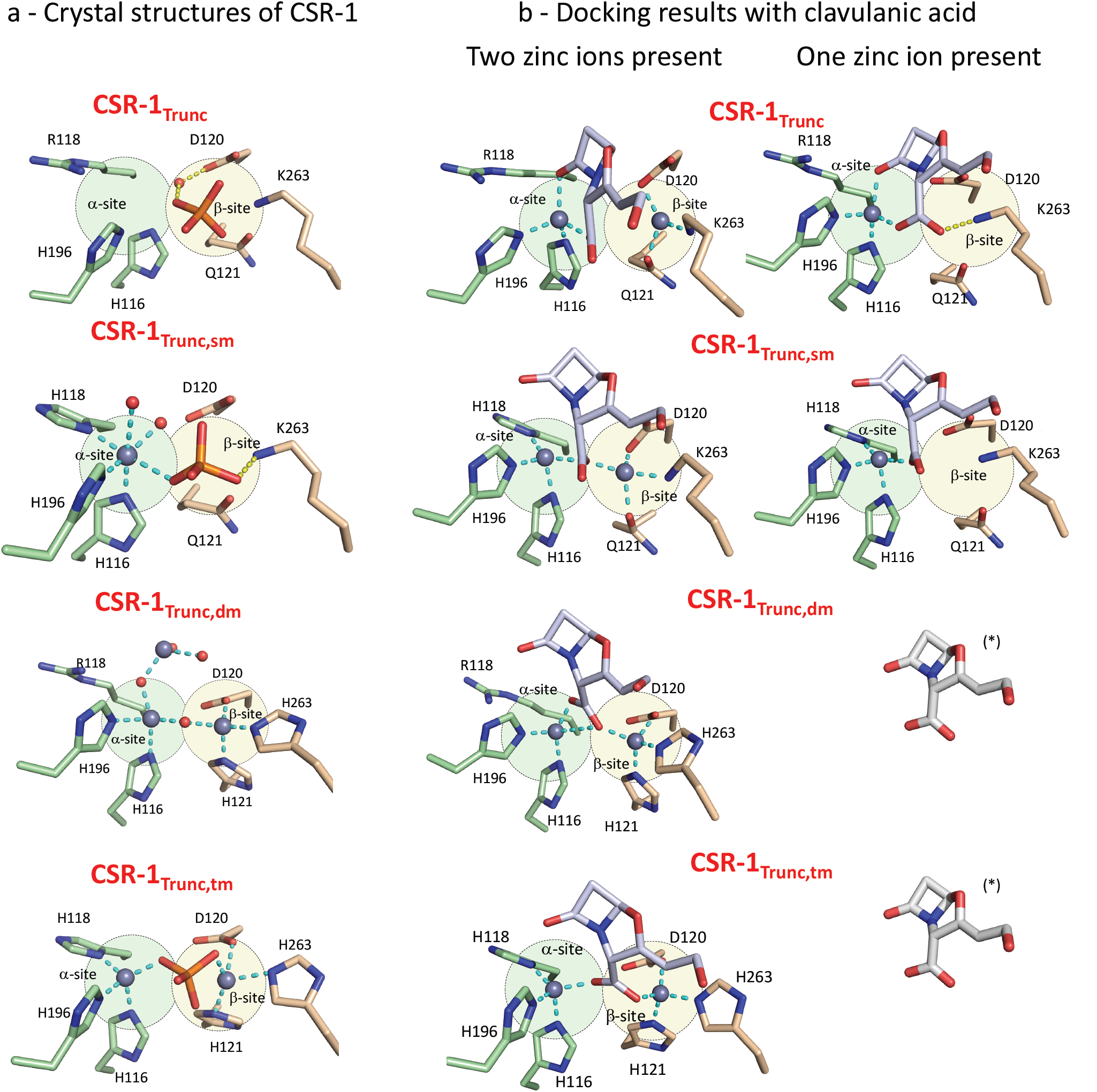
Structural analysis of CSR-1 variants and their interaction with the inhibitor clavulanic acid. **a**. Crystal structures of the active sites of the B3-RQK enzyme CSR-1 and its mutants highlighting the α-(green circle) and β- (yellow circle) metal binding sites. **b**. Predicted docking of clavulanic acid to CSR-1 variants in the presence of one or two Zn^+2^ metal ions. The predicted binding of the bi-metallic forms of CSR-1 and its mutants are unlikely to represent the inhibited form of this enzyme because there is no difference in the four poses (*see main text*). In the presence of one metal ion, however, clavulanic acid(*) is predicted to only form a stable enzyme-inhibitor complex with CSR-1 and its single mutant, consistent with the experimental inhibition data (**Supplemental Table 2**).

### CSR-1 mutations increase antibiotic resistance and reverse sensitivity to clavulanic acid

The increasing Zn^2+^ binding capacity of CSR-1_trunc_ as a function of introduced mutations was paralleled by enhanced antibiotic resistance and increased catalytic activity, including loss of sensitivity to carbenicillin, cefuroxime, meropenem and clavulanic acid in the double and triple mutants (**Table 2**). The effect of the single mutation in the α-site (*i*.*e*. R118H) on the catalytic properties is modest; CSR-1_trunc,sm_ displays a slightly enhanced catalytic activity towards most of the substrates tested, but this positive effect is largely offset by reduced substrate binding. The effect of the double and triple mutations in CSR-1_trunc_, however, is striking. A marked increase in catalytic efficiency was observed against most antibiotics accompanied by an increase in *ex vivo* resistance to both antibiotics (aggregate resistance score 8.5 *vs* 5 for CSR-1_trunc_) and clavulanic acid (**Table 2**). This indicates that the mutations in the β-site are primarily responsible for the loss of catalytic activity, antibiotic resistance and atypical sensitivity to clavulanic acid. To explore this further, we performed *in silico* docking calculations. Calculations based on the bimetallic enzyme predicted that the carboxylate group of clavulanic acid binds to both metal ions and forms hydrogen bonds with Ser214, Asn254 and Arg257 in the substrate binding pocket of CSR-1_trunc_ (**Figure 4b**). Once the calculation was performed with single metal occupancy in the *α*-site (*i*.*e*. the higher metal affinity site), we observed a stable enzyme-inhibitor complex only in CSR-1_trunc_ and CSR-1_trunc,sm_, indicating the ability of clavulanic acid to displace the β-Zn^2+^ from its low affinity binding site in these two variants of CSR-1. We therefore infer that low metal binding affinity in the β-site is necessary for the observed inhibition by clavulanic acid. Hydrogen bonding to Lys263 may provide additional stabilization of the enzyme-clavulanic acid complex in monometallic CSR-1_trunc_ and CSR-1_trunc,sm_, as this interaction cannot be formed in the double and triple mutants, in which His263 strongly interacts with Zn^2+^ bound in the β site (**Figure 4b**).

**Table 2.**
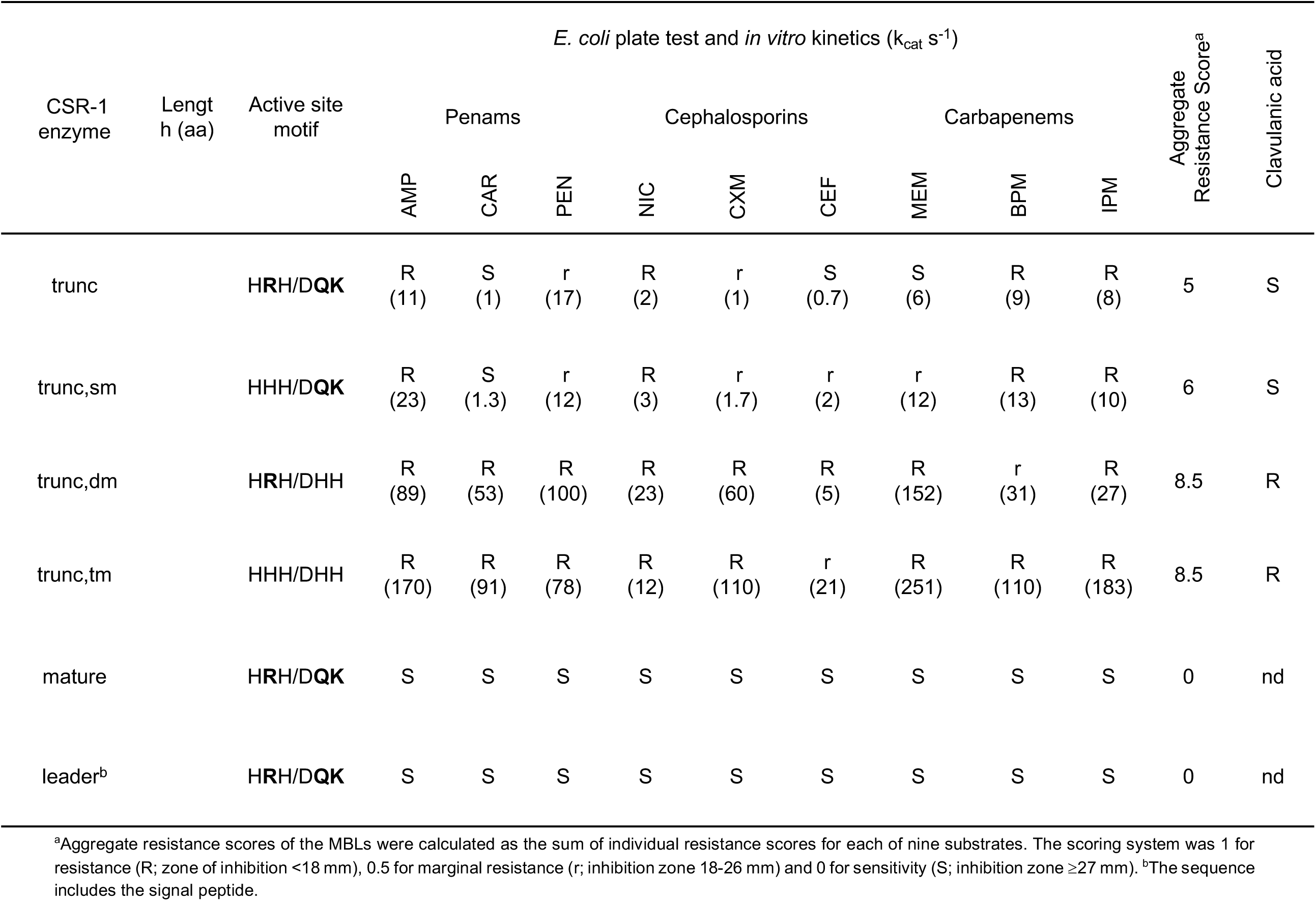
*In vitro* kinetic data and *ex vivo* plate tests for CSR-1 and its mutants.

In summary, despite being phylogenetically unrelated to their counterparts from the B1 (HHH/DCH) and B2 (NHH/DCH) subgroups, canonical B3 MBLs have a similar active site motif (HHH/DHH), with similar catalytic properties. Indeed, B3 MBLs such as the well-studied L1 and AIM-1 enzymes are as efficient as B1 and B2 MBLs in inactivating a broad range of β-lactam antibiotics, and are not inhibited by clavulanic acid (20, 23, 24). Through a broad phylogenetic analysis of B3 MBL enzymes detected in the rapidly expanding microbial genome database (41), we identified B3-like MBLs in a wide range of hosts and habitats (**Fig. 2**), and confirmed three common active site variants of the canonical B3 MBL motif; B3-E (**E**HH/DHH), B3-Q (**Q**HH/DHH) and B3-RQK (H**R**H/D**QK**; **Fig. 1**). B3-E and B3-Q have similar properties to canonical B3 enzymes, however, B3-RQK is remarkable for a number of reasons; i) weak coordination of Zn^2+^ cofactors, ii) is the only B3 with amino acid substitutions in the *β*-site, iii) the N-terminus occludes the active site, iv) enzymatic activity and *ex vivo* conferral of antibiotic resistance are reduced and only present if the N-terminus is truncated, and v) is sensitive to clavulanic acid (**Table 1, Figures 3 & 4**). Mutation of the B3-RQK active site to restore the canonical B3 motif resulted in increased metal binding and catalytic activity, *ex vivo* resistance to a wider range of *β*-lactam antibiotics, and loss of sensitivity to clavulanic acid (**Supplemental Table S2**). This can be primarily attributed to the restoration of the canonical histidine in position 263 of the MBL *β*-site, which greatly enhances the metal ion affinity and prevents binding of clavulanic acid to B3 MBLs and their B3-E and B3-Q variants. B1 and B2 MBLs are largely unaffected by clavulanic acid, with the notable exception of the B1 MBL SPM-1 (48). However, in the present study we provide the first insight into the mechanism that enables clavulanic acid to inhibit an MBL by displacing the catalytically essential Zn^2+^ ion bound in the β site of the active center. Although it is not yet known how clavulanic acid inhibits SPM-1, the presence of the canonical histidine in position 263 in this enzyme indicates that clavulanic acid may be modified to enhance its interaction with His263, thus outcompeting the Zn^2+^ in the β site, and thereby increasing the therapeutic range of this widely used antibiotic resistance drug.

## Methods

### Phylogenetic Analysis

Putative protein orthologues of the B3 family of MBLs were identified from the Genome Taxonomy Database using GeneTreeTK (version 0.0.11; https://github.com/dparks1134/GeneTreeTk). Sequences were manually curated, then aligned with MAFFT. Model parameters for phylogenetic inference were evaluated using ModelFinder in IQ-Tree, selecting the optimal model according to the Bayesian Information Criterion. Detailed experimental procedures are described in SI Appendix.

### *Ex vivo* whole cell plate assays

Genes were cloned into pET27b(+) and transformed into *E. coli* BL21(DE3) cells. Transformed bacterial suspensions (0.5 McFarland units) were plated onto Mueller-Hinton (MH) agar. Antibiotic resistance tests were similarly performed with diffusion discs containing penicillin G, carbenicillin, cephalothin, nitrocefin, cefuroxime, meropenem, imipenem and biapenem. Detailed experimental procedures are described in SI Appendix.

### Protein expression and purification

CSR-1 and its mutants were transformed into chemically competent *E. coli* LEMO BL21 (DE3) cells by heat shock. The cells were inoculated into TB medium and grown at 37°C for 48 hours. The protein was purified using a SP Sepharose column in 20 mM HEPES, pH 7.5, and 0.15 mM of ZnCl_2_, and the protein was eluted with a gradient of 1 M of NaCl. Detailed experimental procedures are described in SI Appendix.

### Crystallization, X-ray diffraction data collection and refinement

Crystals of CSR-1 and its mutants were grown at 18°C by hanging-drop vapor diffusion using a 24-well pre-greased plate (Hampton Research) with drops containing 1 µL of the protein solution and 1 µL of precipitant buffer comprising 0.49 M of NaHSO_4_, 0.91 M of KHSO_4_ and 50 % w/v PEG 400. Diffraction data were collected from cryoprotected crystals on beamline MX-2 at the Australian Synchrotron (Melbourne) using BLU-ICE. Detailed experimental procedures are described in SI Appendix.

### Molecular docking and QM/MM calculations

For the theoretical studies, the missing Zn^2+^ ions in CSR-1_trunc_ and CSR-1_trunc,sm_ were manually placed in the binding site using CSR-1_trunc,tm_ as a template. All water molecules were removed, as well as the artefactual third Zn^2+^ in CSR-1_trunc,dm_ (**Figure 4a**). Proteins were protonated using the program *tleap* in the AmberTools16 software package. Selected poses from the initial molecular docking calculations were further optimized using a QM/MM-based potential according to Marion *et al*. The QM/MM calculations were performed using the ChemShell suite with the DL_POLY module for the MM-calculations interfacing with Turbomole version 7.1 for the QM-calculations. Detailed experimental procedures are described in SI Appendix.

## Acknowledgements

This research was supported by Project Grants from the NH&MRC (APP1084778) and Australian Research Council (DP150104358), a Future Fellowship (FT120100694) awarded to GS, and a Laureate Fellowship (FL150100038) awarded to PH. NM was supported by a Science Foundation Ireland - President of Ireland Young Researcher Award (SFI-PIYRA). IA and OM were supported by the Deutsche Forschungsgemeinschaft (SFB 749, project C08; and CIPSM).

## Author Contribution

MMP, NM, RPM and GS devised the study. MMP, NM and LW performed the experimental work. MMP, DWW, OM, IA and LWG performed the bioinformatic analysis. MMP, PH and GS wrote the manuscript, and all authors contributed edits to the final version.

## Additional information

None

