## supplemental material for "Broad spectrum antibiotic-degrading metallo-β-lactamases are phylogenetically diverse and widespread in the environment"

### Methods

#### Phylogenetic Analysis

Putative protein orthologues of the B3 family of MBLs were identified from the Genome Taxonomy Database using GeneTreeTK (version 0.0.11; https://github.com/dparks1134/GeneTreeTk). Sequences were manually curated, then aligned with MAFFT (1). Columns representing six residues critical for metal ion binding were manually identified in the L1 alignment (His105, His107 and His181 for the α site, and Asp109, His110 and His246 for the β site), and proteins categorized according to their motif (B3: HHH/DHH, B3-RQK: HRH/DQK, B3-Q: QHH/DHH and B3-E: EHH/DHH for their α/β metal binding sites, respectively). For phylogenetic inference, sequences were dereplicated into clusters of proteins sharing at least 70% amino acid identity using usearch (v8.1) (2). A total of 673 representative protein sequences (518 B3, 77 B3-Q, 35 B3-E and 43 B3-RQK) were aligned using MAFFT and columns filtered using TrimAl (1). Model parameters for phylogenetic inference were evaluated using ModelFinder (3) in IQ-Tree (4), selecting the optimal model according to the Bayesian Information Criterion. Inference was then performed in IQ-Tree using the WAG+F+R10 model of amino acid substitution and 100 bootstrap re-samplings to assess node support. The resulting tree was visualized and annotated with genome metadata using the Interactive Tree of Life resource (5).

#### *Ex vivo* whole cell plate assays

All genes listed in **Table 1**, with the exception of FEZ-1, were synthesized by General Biosystems. Genes were cloned into pET27b(+) and transformed into *E. coli* BL21(DE3) cells. Transformed bacterial suspensions (0.5 McFarland units) were plated onto Mueller-Hinton (MH) agar. Antibiotic resistance tests were similarly performed with diffusion discs containing penicillin G, carbenicillin, cephalothin, nitrocefin, cefuroxime, meropenem, imipenem and biapenem. Plates were incubated at 35°C for 18 hours to allow bacterial growth, and zones of clearing around strips and/or discs were measured with a caliper against a dark background. Tests were conducted in quadruplicate, and the inhibition zone measurements averaged. *E. coli* cells with pET22b(+) plasmids lacking inserts were used as negative controls. Strip and disc tests were conducted according to the European Society of Clinical Microbiology and Infectious Diseases (EUCAST) standard for antimicrobial susceptibility testing.

#### Protein expression and purification

CSR-1, CSR-$1_{trunc}$, CSR-$1_{trunc,sm}$, CSR-$1_{trunc,dm}$ and CSR-$1_{trunc,tm}$ constructs were transformed into chemically competent *E. coli* LEMO BL21 (DE3) cells by heat shock. A single colony was inoculated into 20 mL Luria-Bertani medium supplemented with 50 µg/ml of kanamycin and 30 µg/ml of chloramphenicol, and the cultures were grown overnight at 37°C and then inoculated into Terrific Broth medium (1 L) and grown at 37°C until an optical density ($OD_{600}$) of 0.4-0.6 was reached. The temperature was then reduced to 18°C and liquid media incubated for an additional 48 hours. Subsequently, the cells were harvested by centrifugation at 4°C and resuspended in 20 mM HEPES, pH 7.5, and 0.15 mM of $ZnCl_2$, protease inhibitor cocktail, 1 mg/mL lysozyme, 20 μg/mL of DNAse and $MgCl_2$ (5mM). Cells were disrupted by sonication and the supernatant was loaded into a 25 mL SP Sepharose FF column (GE Healthcare) equilibrated with 20 mM HEPES, pH 7.5, and 0.15 mM of $ZnCl_2$. Expressed target proteins were eluted using a gradient of 1M NaCl in 20 column volumes.

**Crystallization, X-ray diffraction data collection and refinement**

Crystals of the expressed proteins were grown at 18°C by hanging-drop vapor diffusion using a 24-well pre-greased plate (Hampton Research) with drops containing 1 μL of the protein solution and 1 μL of precipitant buffer comprising 0.49 M of $NaHSO_4$, 0.91 M of $KHSO_4$ and 50 % w/v PEG 400. Diamond like crystals were observed after five days of incubation. Harvested crystals were cryoprotected in precipitant buffer containing 20% glycerol. Diffraction data were collected from cryoprotected crystals on beamline MX-2 at the Australian Synchrotron (Melbourne) using BLU-ICE (6). Diffraction data were integrated, scaled and merged using the software HKL-2000 (7). Refinement and model building were carried out using PHENIX 1.8.4 (8) and COOT 0.7 (9), respectively. The CSR-1 and CSR-$1_{trunc}$ structures were determined using the previously published coordinates for L1 (PDB 1SML) (10). The mutants CSR-$1_{trunc,sm}$, CSR-$1_{trunc,dm}$ and CSR-$1_{trunc,tm}$ were then solved by molecular replacement using the CSR-1 structure as a search model. All atoms were subsequently refined with anisotropic B-factors; most hydrogen atoms were fitted as riding models. Relevant crystallographic data and refinement statistics are summarized in **Supplemental Table 5**.

**Enzyme Kinetics**

All kinetic measurements were performed on a Cary 60 Bio Varian UV-Vis spectrophotometer. In a standard assay, reactions were initiated by the addition of the enzyme and monitored for 60 s at 25°C. The Michaelis-Menten (11) parameters were determined by nonlinear fitting using GraphPad Prism 7. The hydrolysis of ampicillin, penicillin G, carbenicillin, nitrocefin, cefoxitin, cefuroxime, meropenem, biapenem, imipenem and nitrocefin were monitored in 50 mM TrisHCl (pH 8.5) by detecting the hydrolysis of the substrate. Inhibition measurements were obtained by measuring the activity of the enzyme with ampicillin and analyzing the effect of the increase in clavulanic acid concentration on the enzyme activity. The data were calculated using the mixed-inhibition equation (11).

**Metal binding studies**

The metal-free apo forms of the CSR-1 enzyme and its mutants were obtained by incubating approximately 3 mg of protein in a 3 mL solution containing 10 mM of EDTA in 20 mM HEPES buffer (pH 7.0 at 4°C). After 24 hours, protein samples were separated from the chelating solution using an Econo-Pac 10DG gel filtration column equilibrated with fresh HEPES buffer treated with Chelex resin to remove any residual metal ions. Atomic absorption spectroscopy (AAS) was used to confirm the absence of metal ions in the final protein solutions. $Zn^{2+}$ ions were then titrated back into the protein solutions and the heat release measured by isothermal titration calorimetry (ITC) at 25°C using a Nano ITC from TA instruments. The concentration of the metal ion solution was verified against standardized EDTA solutions and the results were compared against AAS data. At least four sets of data were collected per protein and the average calculated. The ITC data were fitted using TRIOS v4.4.0 software from TA instruments. The software uses a nonlinear algorithm to minimize $\chi^2$ values to fit the experimental data to equations derived from equilibrium-binding models simulating either one or two independent binding sites (**Supplemental Table 4**).

**Molecular docking and QM/MM calculations**

For the theoretical studies, the missing $Zn^{2+}$ ions in CSR-$1_{trunc}$ and CSR-$1_{trunc,sm}$ were manually placed in the binding site using CSR-$1_{trunc,tm}$ as a template. All water molecules were removed, as well as the artefactual third $Zn^{2+}$ in CSR-$1_{trunc,dm}$ (**Figure 4a**). Proteins were protonated using the program *tleap* in the AmberTools16 software package (12). Metal-ligating histidine residues were protonated at the nitrogen not involved in metal coordination. Lys263 in the β-site was considered in its neutral form such that it can ligate a $Zn^{2+}$ ion. Molecular docking of clavulanic acid was performed with FlexX (13) within the LeadIT platform version 2.3.2 (https://www.biosolveit.de/LeadIT/). All residues within 10 Å of the geometric center of the two metal ions were considered as binding site residues. The 'Enthalpy and Entropy' (*i.e.* Hybrid) approach was used for initial base placement and the clash factor and maximum allowed overlap volume were set to 0.6 and 3.5 $Å^3$, respectively.

Selected poses from the initial molecular docking calculations were further optimized using a QM/MM-based potential according to Marion *et al.* (14) The QM/MM calculations were performed using the ChemShell suite (15, 16) with the DL_POLY module for the MM-calculations interfacing with Turbomole (17) version 7.1 for the QM-calculations. The MM-region was described applying the parameters from the Amber ff14SB (18) force field. The QM-region comprised the side chains of Asn142, His196, Leu197, Asp208, Ser214, Tyr229, Asn262, Lys263, Glu265 and Arg266, and the loop from Asn114 to Gln121, as shown in **Supplemental Figure 4**. The QM/MM boundary was located at the nonpolar $C_\alpha$-$C_\beta$ bond of the single side chains and at the N-$C_\alpha$ and $C_\alpha$-C' bond of the backbone, respectively. The QM-calculations were performed at the DFT level together with the D3 dispersion correction (19). The TPSS meta-GGA (20) functional was applied using RI-J approximation (21) on a multigrid m4 (22). The SCF convergence criterion was set to $10^{-7}$ au. The def2-TZVP (22, 23) basis set was used for the $Zn^{2+}$ ions while the remainder of the QM-region was described with the def2-SVP (22, 24) basis set. An electrostatic embedding scheme was applied to describe the QM/MM boundary, in which link atoms are placed at the boundary and the charges are shifted away and replaced by a dipole on the recipient atom, as implemented in the polarized coupling scheme (*shift)* in ChemShell. The geometry optimization of all atoms in the QM-region and all direct neighboring MM-residues was performed using the DL-FIND (25) optimizer.

### Supplementary Tables

#### Supplementary Table 1. Number of copies found in the different species.

| Genome | Genome (NCBI) | Tip label | B3 | B3-E | B3-Q | B3-RQK | Total |
|---|---|---|---|---|---|---|---|
| GB_GCA_001767525.1 | GCA_001767525.1 | Acidobacteria bacterium RIFCSPLOWO2_12_FULL_68_19 | 5 | 0 | 0 | 0 | 5 |
| GB_GCA_001767195.1 | GCA_001767195.1 | Acidobacteria bacterium RIFCSPLOWO2_02_FULL_67_21 | 3 | 0 | 0 | 0 | 3 |
| GB_GCA_001898925.1 | GCA_001898925.1 | Novosphingobium sp. 63-713 | 1 | 2 | 0 | 0 | 3 |
| GB_GCA_001919455.1 | GCA_001919455.1 | Acidobacteria bacterium 13_1_40CM_2_56_11 | 3 | 0 | 0 | 0 | 3 |
| RS_GCF_000379645.1 | GCF_000379645.1 | Actinoplanes globisporus DSM 43857 | 3 | 0 | 0 | 0 | 3 |
| RS_GCF_001485665.1 | GCF_001485665.1 | Janthinobacterium sp. CG23_2 | 3 | 0 | 0 | 0 | 3 |
| RS_GCF_001865675.1 | GCF_001865675.1 | Janthinobacterium sp. 1_2014MBL_MicDiv | 2 | 1 | 0 | 0 | 3 |
| GB_GCA_000974665.1 | GCA_000961635.1 | alpha proteobacterium U9-1i | 2 | 0 | 0 | 0 | 2 |
| GB_GCA_001464575.1 | GCA_001464575.1 | Acidobacteria bacterium Ga0077553 | 1 | 0 | 1 | 0 | 2 |
| GB_GCA_001618865.1 | GCA_001618865.1 | Acidobacteria bacterium DSM 100886 | 0 | 1 | 1 | 0 | 2 |
| GB_GCA_001899605.1 | GCA_001899605.1 | Sphingobacteriales bacterium 40-81 | 1 | 0 | 1 | 0 | 2 |
| GB_GCA_001899685.1 | GCA_001899685.1 | Sphingobacteriales bacterium 44-15 | 1 | 0 | 1 | 0 | 2 |
| GB_GCA_900103265.1 | GCA_900103265.1 | Sphingomonas sp. NFR15 | 1 | 1 | 0 | 0 | 2 |
| GB_GCA_900114495.1 | GCA_900114495.1 | Dyella sp. OK004 | 1 | 1 | 0 | 0 | 2 |
| RS_GCF_000014905.1 | GCF_000014905.1 | Candidatus Solibacter usitatus Ellin6076 | 1 | 0 | 1 | 0 | 2 |
| RS_GCF_000017265.1 | GCF_000017265.1 | Phenylobacterium zucineum HLK1 | 2 | 0 | 0 | 0 | 2 |
| RS_GCF_000204015.1 | GCF_000204015.1 | Asticcacaulis biprosthecum C19 | 2 | 0 | 0 | 0 | 2 |
| RS_GCF_000242815.1 | GCF_000242815.1 | Janthinobacterium lividum PAMC 25724 | 1 | 1 | 0 | 0 | 2 |
| RS_GCF_000253255.1 | GCF_000253255.1 | MIM-1 | 2 | 0 | 0 | 0 | 2 |
| RS_GCF_000281975.1 | GCF_000281975.1 | Novosphingobium sp. AP12 | 1 | 1 | 0 | 0 | 2 |
| RS_GCF_000374905.1 | GCF_000374905.1 | Arthrobacter sp. 162MFSha1.1 | 2 | 0 | 0 | 0 | 2 |
| RS_GCF_000381565.1 | GCF_000381565.1 | Oxalobacteraceae bacterium AB_14 | 2 | 0 | 0 | 0 | 2 |
| RS_GCF_000382885.1 | GCF_000382885.1 | Sphingobium japonicum BiD32 | 2 | 0 | 0 | 0 | 2 |
| RS_GCF_000447205.1 | GCF_000447205.1 | Sphingobium ummariense RL-3 | 1 | 1 | 0 | 0 | 2 |
| RS_GCF_000473995.1 | GCF_000473995.1 | Clostridium saccharobutylicum DSM 13864 | 2 | 0 | 0 | 0 | 2 |
| RS_GCF_000632105.1 | GCF_000632105.1 | Novosphingobium resinovorum | 2 | 0 | 0 | 0 | 2 |

| | | | | | | | |
|---|---|---|---|---|---|---|---|
| RS_GCF_000744465.1 | GCF_000744465.1 | Caulobacter henricii | 2 | 0 | 0 | 0 | 2 |
| RS_GCF_000763945.1 | GCF_000763945.1 | Sphingopyxis sp. MWB1 | 1 | 1 | 0 | 0 | 2 |
| RS_GCF_000813185.1 | GCF_000813185.1 | Novosphingobium sp. MBES04 | 1 | 1 | 0 | 0 | 2 |
| RS_GCF_000829715.2 | GCF_000829715.2 | Streptomyces sp. NBRC 110027 | 2 | 0 | 0 | 0 | 2 |
| RS_GCF_001298105.1 | GCF_001298105.1 | Novosphingobium sp. ST904 | 1 | 1 | 0 | 0 | 2 |
| RS_GCF_001421325.1 | GCF_001421325.1 | Novosphingobium sp. Leaf2 | 1 | 1 | 0 | 0 | 2 |
| RS_GCF_001426905.1 | GCF_001426905.1 | Caulobacter sp. Root1455 | 2 | 0 | 0 | 0 | 2 |
| RS_GCF_001427665.1 | GCF_001427665.1 | Caulobacter sp. Root655 | 1 | 1 | 0 | 0 | 2 |
| RS_GCF_001449105.1 | GCF_001449105.1 | Caulobacter vibrioides | 1 | 1 | 0 | 0 | 2 |
| RS_GCF_001591025.1 | GCF_001591025.1 | Sphingomonas soli NBRC 100801 | 2 | 0 | 0 | 0 | 2 |
| RS_GCF_001591045.1 | GCF_001591045.1 | Sphingopyxis granuli NBRC 100800 | 2 | 0 | 0 | 0 | 2 |
| RS_GCF_001598415.1 | GCF_001598415.1 | Sphingomonas mali NBRC 15500 | 1 | 1 | 0 | 0 | 2 |
| RS_GCF_001717955.1 | GCF_001717955.1 | Sphingomonas panacis | 1 | 1 | 0 | 0 | 2 |
| RS_GCF_002001165.1 | GCF_002001165.1 | Rhodanobacter sp. C05 | 2 | 0 | 0 | 0 | 2 |
| RS_GCF_002007115.1 | GCF_002007115.1 | Massilia sp. KIM | 2 | 0 | 0 | 0 | 2 |
| RS_GCF_900011245.1 | GCF_900011245.1 | Bradyrhizobium sp. | 2 | 0 | 0 | 0 | 2 |
| RS_GCF_900156165.1 | GCF_900156165.1 | Pseudacidovorax sp. RU35E | 2 | 0 | 0 | 0 | 2 |
| RS_GCF_900177785.1 | GCF_900177785.1 | Cedecea sp. NFIX57 | 1 | 0 | 0 | 1 | 2 |
| U_66028 | SAMN08018157 | Gammaproteobacteria bacterium UBA8143 | 1 | 0 | 1 | 0 | 2 |
| U_66329 | GCA_002311835.1 | Proteobacteria bacterium UBA1148 | 0 | 0 | 2 | 0 | 2 |
| U_67505 | SAMN08019550 | Gammaproteobacteria bacterium UBA8658 | 1 | 0 | 1 | 0 | 2 |
| U_67723 | GCA_002336595.1 | Gammaproteobacteria bacterium UBA1979 | 1 | 0 | 1 | 0 | 2 |
| U_68064 | GCA_002328835.1 | Acidobacteria bacterium UBA2161 | 2 | 0 | 0 | 0 | 2 |
| U_69330 | GCA_002347025.1 | Acidobacteria bacterium UBA2990 | 2 | 0 | 0 | 0 | 2 |
| U_69335 | GCA_002346545.1 | Acidobacteria bacterium UBA2994 | 1 | 0 | 1 | 0 | 2 |
| U_72084 | GCA_002387245.1 | Brevundimonas sp. UBA4553 | 1 | 1 | 0 | 0 | 2 |
| U_74706 | SAMN08018207 | Janthinobacterium bacterium UBA11349 | 2 | 0 | 0 | 0 | 2 |
| U_74816 | GCA_002428565.1 | Anaerolineales bacterium UBA6092 | 2 | 0 | 0 | 0 | 2 |
| U_75494 | GCA_002434915.1 | Proteobacteria bacterium UBA6522 | 0 | 0 | 2 | 0 | 2 |
| U_77156 | GCA_002477465.1 | Gammaproteobacteria bacterium UBA7449 | 1 | 0 | 1 | 0 | 2 |
| U_77191 | GCA_002478575.1 | Sphingomonadales bacterium UBA7473 | 1 | 1 | 0 | 0 | 2 |

### Supplementary Table 2. *Ex vivo* sensitivity disc test.

| Class | Enzyme | Host species | *Sensitivity Tests – results in millimetres (mm)* | | | | | | | | | | |
|---|---|---|---|---|---|---|---|---|---|---|---|---|---|
| | | | Penams | | | Cephalosporins | | | Carbapenems | | | Clavulanic Acid Inhibition | |
| | | | AMP | CAR | PEN | NIC | CXM | CEF | MEM | BPM | IPM | [a]AML | [b]AMLC |
| B3 | L1 | *Stenotrophomonas maltophilia* | 5±1 | 15±5 | 14±7 | 11±2 | 22±7 | 12±7 | 22±3 | 20±1 | 10±7 | 5±7 | 5±5 |
| | AIM-1 | *Pseudomonas aeruginosa* | 5.0±1 | 10±1 | 15±3 | 12±1 | 13±3 | 21±4 | 36±7 | 12±7 | 10±2 | 7±5 | 6±4 |
| | MIM-1 | *Novosphingobium pentaromativorans* | 18±4 | 15±7 | 21±3 | 10±7 | 5±7 | 34±7 | 40±6 | 15±5 | 12±5 | 5±3 | 5±2 |
| | MIM-2 | *Simiduia agarivorans* | 18±2 | 10±4 | 7±4 | 19±3 | 5±7 | 24±2 | 42±5 | 12±3 | 12±7 | 8±4 | 7±3 |
| B3-RQK | SPR-1$_{Trunc}$ | *Serratia proteamaculans* | 5±2 | 42±3 | 20±2 | 12±3 | 22±2 | 45±3 | 44±2 | 24±2 | 16±1 | 6±4 | 16±4 |
| | CSR-1$_{Trunc}$ | *Cronobacter sakasakii* | 10±2 | 44±3 | 18±4 | 15±1 | 21±3 | 43±2 | 47±2 | 15±3 | 12±2 | 5±7 | 17±2 |
| | SER-1$_{Trunc}$ | *Salmonella enterica* | 5±1 | 47±3 | 19±4 | 12±3 | 9±4 | 44±4 | 45±1 | 14±2 | 18±3 | 7±3 | 28±4 |
| B3-Q | GOB-1 | *Elizabethkingia meningoseptica* | 8±4 | 19±2 | 5±2 | 6±4 | 8±2 | 18±3 | 21±4 | 47±3 | 5±4 | 5±4 | 5±1 |
| | CMQ-1 | *Chryseobacterium molle* | 7±4 | 41±3 | 18±2 | 10±7 | 9±7 | 27±4 | 30±4 | 30±7 | 5±4 | 5±3 | 4±8 |
| | SIQ-1 | *Sphingomonas indica* | 10±4 | 5±2 | 7±4 | 6±3 | 5±2 | 9±3 | 33±6 | 30±8 | 5±5 | 6±5 | 5±9 |
| B3-E | SIE-1 | *Sphingobium indicum* | 15±2 | 10±6 | 7±4 | 5±5 | 5±2 | 43±7 | 40±3 | 35±4 | 5±1 | 9±4 | 9±2 |
| | SSE-1 | *Sphingopyxis sp.* | 5±3 | 5±7 | 10±2 | 9±5 | 5±2 | 18±2 | 33±6 | 41±4 | 5±4 | 7±4 | 6±1 |

[a] Amoxycillin
[b] Amoxycillin and clavulanic acid

**Supplementary Table 3. Steady-state kinetic parameters for CSR-1$_{trunc}$, CSR-1$_{trunc,sm}$, CSR-1$_{trunc,dm}$ and CSR-1$_{trunc,tm}$.**

| | $k_{cat}$ ($s^{-1}$) | | | |
|---|---|---|---|---|
| **Substrate** | **CSR-1$_{trunc}$** | **CSR-1$_{trunc,sm}$** | **CSR-1$_{trunc,dm}$** | **CSR-1$_{trunc,tm}$** |
| Ampicillin | 11±2 | 23±8 | 89±7 | 170±9 |
| Penicillin G | 17±3 | 12±7 | 100±9 | 78±2 |
| Carbenicillin | 1±0.7 | 1.3±1 | 53±2 | 91±7 |
| Cefuroxime | 1±0.9 | 1.7±0.9 | 60±3 | 110±7 |
| Cefoxitin | 0.7±0.6 | 2±0.7 | 5±1 | 21±8 |
| Nitrocefin | 2±0.3 | 3±1 | 23±7 | 12±1 |
| Imipenem | 8±1 | 10±2 | 31±2 | 183±13 |
| Meropenem | 6±3 | 12±2 | 152±9 | 251±3 |
| Biopenem | 9±2 | 13±5 | 27±7 | 110±9 |

**Supplementary Table 4. ITC parameters collected for CSR-1$_{trunc}$ and its mutants.**

| | **CSR-1$_{trunc}$** | **CSR-1$_{trunc,sm}$** | **CSR-1$_{trunc,dm}$** | **CSR-1$_{trunc,tm}$** |
|---|---|---|---|---|
| Model[(a)] | Two Independent sites | Two Independent sites | One Independent Site | Two Independent sites |
| $N_1$ | 0.95±0.09 | 0.98±0.07 | 1.99±0.1 | 0.99±0.9 |
| $K_{d1}$ (μM) | 2±0.7 | 0.1±0.07 | 0.14±0.07 | 0.06±0.02 |
| $N_2$ | 0.87±0.06 | 1.2±0.06 | - | 0.83±0.1 |
| $K_{d2}$ (μM) | 150±12 | 180±27 | - | 0.4±0.1 |

[a]Mathematical model applied to fit the experimental data. N is the stoichiometry of an interaction measured by ITC, and $K_d$ is the dissociation constant.

**Supplementary Table 5. Crystallography parameters for the CSM-1 and its mutants.**

| | | | | | |
|---|---|---|---|---|---|
| | | | **Data collection parameters** | | |
| | CSM-1 | CSM-$1_{trunc}$ | CSM-$1_{trunc,sm}$ | CSM-$1_{trunc,dm}$ | CSM-$1_{trunc,tm}$ |
| Resolution Range (Å) | 42.29-1.99 (2.04-1.99) [a] | 35.16-2.16 (2.22-2.16) | 36.90-1.10 (1.12-1.10) | 43.75-2.07 (2.52-2.07) | 43.77-1.48 (1.50-1.48) |
| Observations (I>σ(I)) | 137538 (8753) | 57051(17897) | 933261 (20130) | 126532(8265) | 276496 (9478) |
| Unique reflections (I>σ(I)) | 18134 (1236) | 17930 (1129) | 86879 (2205) | 19081(1376) | 39943 (1589) |
| Completeness (%) | 99.7 (98.3) | 95.0 (93.0) | 93.9 (48.9) | 98.0(87.8) | 99.0 (81.0) |
| Mean <I/σ(I)> | 12.0 (3.1) | 10.9 (1.3) | 12.7 (1.8) | 18.4(2.1) | 10.1 (2.0) |
| [b]$R_{merge}$ | 0.138 (0.544) | 0.095 (0.701) | 0.094 (1.254) | 0.076(0.689) | 0.158 (1.074) |
| [c]$R_{p.i.m.}$ | 0.053 (0.217) | 0.082 (0.638) | 0.030 (0.417) | 0.048(0.441) | 0.064 (0.455) |
| Multiplicity | 7.6 (7.1) | 3.2 (2.8) | 10.7 (9.1) | 5.0(6.0) | 6.9 (6.0) |
| Space group | *I*4 | $P\ 3_1\ 2\ 1$ | $P\ 2_1\ 2_1\ 2_1$ | $P\ 2_1\ 2_1\ 2_1$ | $P\ 2_1\ 2_1\ 2_1$ |
| Unit cell lengths (Å) | $a = b = 84.58$ $c = 74.98$ | $a = b = 74.41$ $c = 107.47$ | $a = 43.77$ $b = 68.60$ $c = 76.25$ | $a = 53.47$ $b = 74.10$ $c = 76.10$ | $a = 43.77$ $b = 69.15$ $c = 77.52$ |
| Unit cell angles (°) | α = β = γ = 90 | α = β = 90 γ = 120 | α = β = γ = 90 | α = β = γ = 90 | α = β = γ = 90 |
| | | | **Refinement statistics** | | |
| $R_{work}$ | 0.1479 | 0.1934 | 0.1700 | 0.1896 | 0.1510 |
| $R_{free}$ | 0.1808 | 0.2241 | 0.1838 | 0.2202 | 0.1719 |
| rmsd bond lengths (Å) | 0.0025 | 0.0029 | 0.011 | 0.041 | 0.005 |
| rmsd bond angles (°) | 0.58 | 0.579 | 1.65 | 0.549 | 0.85 |
| [d]Clash score | 1.90 | 0.80 | 7.60 | 3.50 | 0.80 |
| | | | **Ramachandran plot statistics** | | |
| Favoured regions | 97.41 | 96.48 | 97.96 | 95.33 | 98.00 |
| Outlier regions | 0.00 | 0.39 | 0.00 | 0.00 | 0.00 |
| Rotamer outliners | 1.45 | 2.55 | 3.55 | 0.51 | 1.03 |
| PDB code | 6DN4 | 6DQ2 | 6DQH | 6NC5 | 6DR8 |

[a]The values in parentheses are for the outer resolution shell. [b]$R_{merge} = \sum_{hkl}\sum_i \left|I_i(hkl) - \left(I(hkl)\right)\right| / \sum_{hkl}\sum_i I_i(hkl)$. [c]$R_{p.i.m.} = \sum_{hkl}\left[\frac{1}{[N(hkl)-1]}\right]^{1/2} \sum_i \left|I_i(hkl) - \left(I(hkl)\right)\right| / \sum_{hkl}\sum_i I_i(hkl)$, where $I_i(hkl)$ is the observed intensity and $I(hkl)$ is the average intensity obtained from multiple observations of symmetry-related reflections. [d] Clashscore is defined as the number of bad overlaps ≥0.4 Å per thousand atoms.

### Supplementary Figures

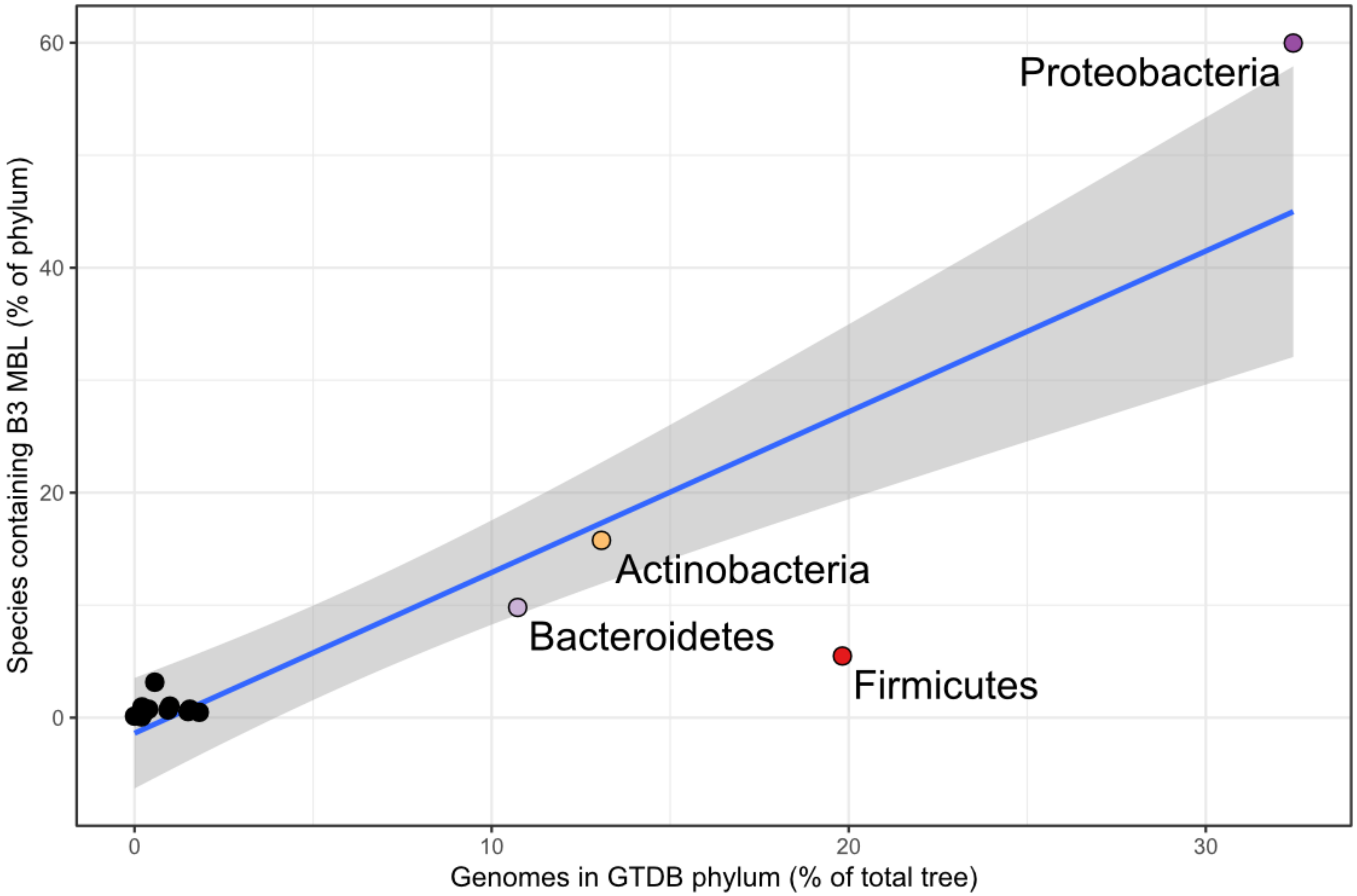


**Supplementary Figure 1. Scatter plot of percentage total genomes *vs* percentage of discovered B3 MBLs in those genomes according to phylum affiliation.** Dominant phyla in the genome database have the highest percentage of B3 MBLs, however, this correlation is not exact as *Proteobacteria* are relatively over-represented and *Firmicutes* relatively under-represented, possibly reflecting greater activity of MBLs on Gram-negative than Gram-positive bacteria.

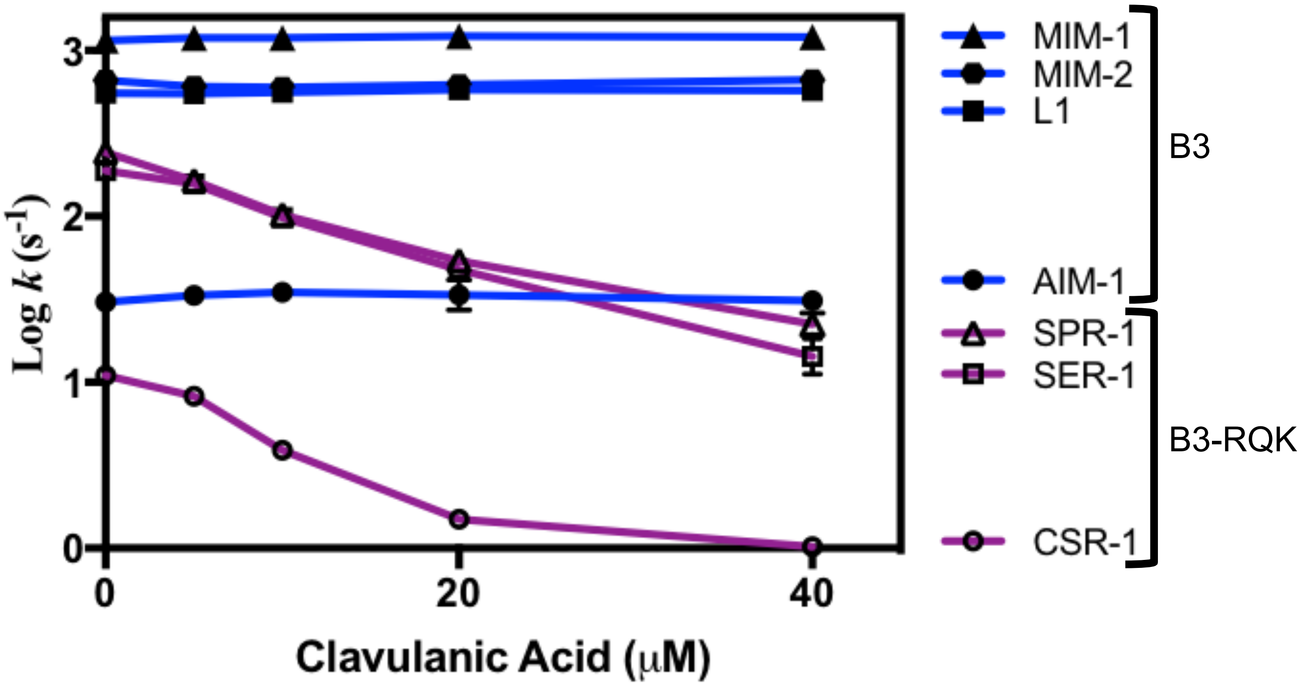


**Supplementary Figure 2. Inhibition of B3 MBLs by clavulanic acid.** The plot shows the effect of different concentrations of clavulanic acid *vs* the logarithm of $k_{cat}$ values of selected B3 MBLs.

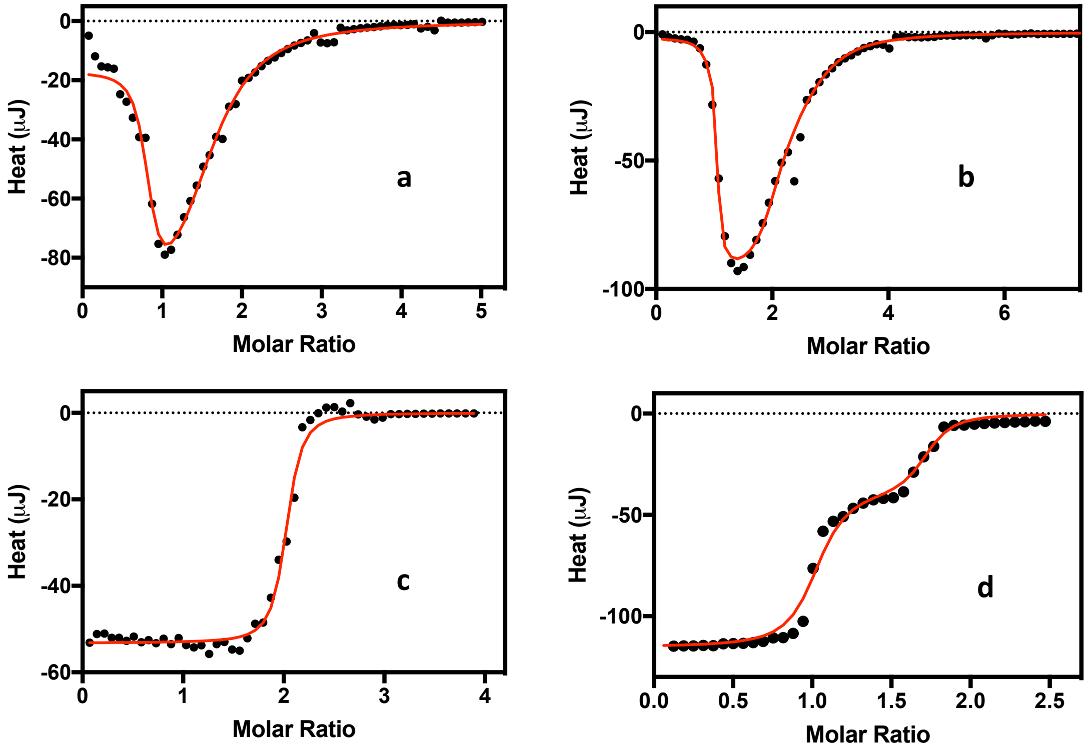


**Supplementary Figure 3. Experimental ITC data collected for the variants of CSR-1 from the subgroup B3-RQK.** Data for CSR-$1_{trunc}$ (a), CSR-$1_{trunc, sm}$ (b), CSR-$1_{trunc,dm}$ (c) and CSR-$1_{trunc,tm}$ (d) were fitted using a nonlinear algorithm to minimise $\chi^2$ values using equations derived from equilibrium-binding models simulating either one or two independent binding sites.

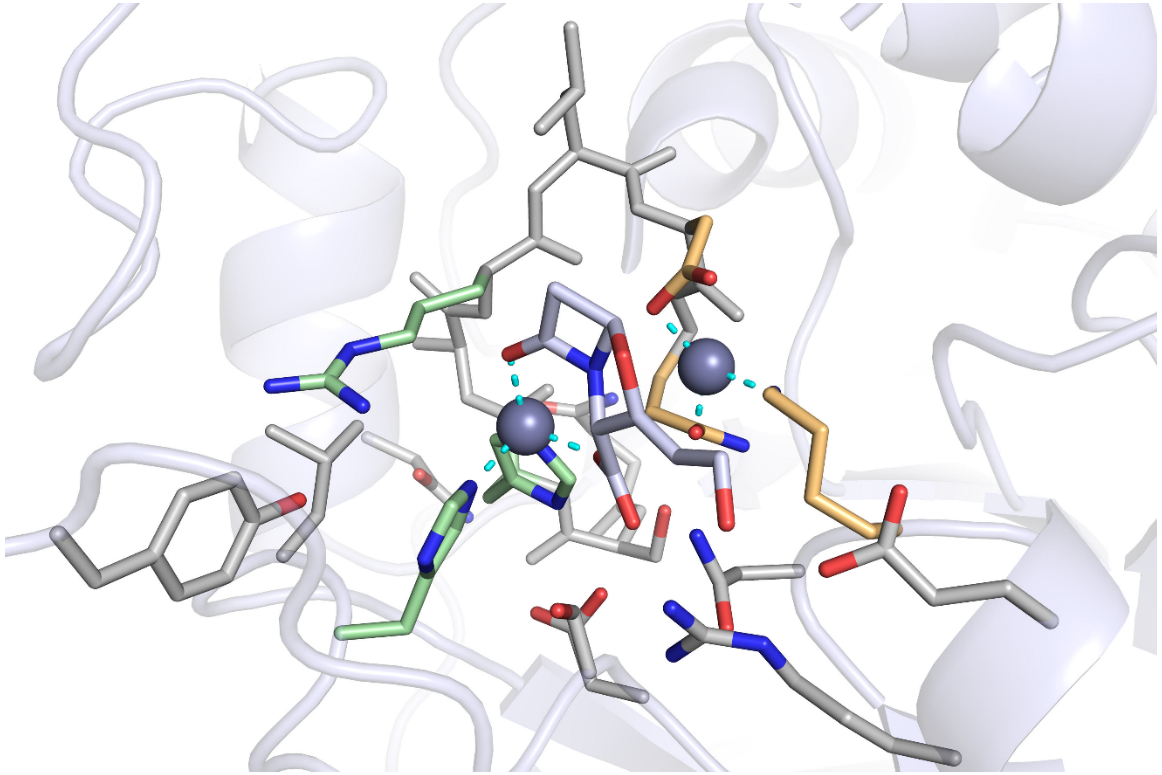

**Supplementary Figure 4. QM region of CSR-1$_{trunc}$, hydrogen atoms are not shown.** $Zn^{2+}$ ions are represented as gray spheres, protein atoms and clavulanic acid as liquorice (carbon atoms of protein and clavulanic acid colored in respective gray and cyan and oxygen red).
